# OmniSplice: detection of non-canonical splicing events from RNA-seq

**DOI:** 10.1101/2025.04.06.647416

**Authors:** Romain Lannes, Rebecca Y. Li, Jaclyn M. Fingerhut, Ryan A. Cummings, Alina D. Salagean, Yukiko M. Yamashita

## Abstract

Splicing generates mature mRNA by removing introns from nascent transcripts and is widely studied using RNA sequencing. However, most RNA-seq analysis pipelines classify RNA-seq reads according to predefined splice-junction structures and discard those that do not conform to such predefined models, potentially obscuring biologically meaningful splicing events. In this study, we developed OmniSplice, a computational framework that captures and analyzes RNA-seq reads that overlap annotated exon ends without assuming predefined splicing architectures. This approach enables systematic detection of non-canonical splicing events that are often overlooked by conventional analyses. Applying OmniSplice to *Drosophila* splicing factor mutants and mouse TDP-43 mutant datasets, we found widespread splicing defects with non-canonical junctions that were not previously recognized, including back-splicing and trans-splicing. Together, these results demonstrate that RNA-seq datasets may contain a substantial reservoir of overlooked splicing information, warranting more comprehensive approaches for analyzing RNA-seq data for splicing events.

## Introduction

Introns are prevalent in eukaryotic genes, and their removal by splicing is essential for the production of mature mRNAs (Nilsen and Graveley, 2010; Rogozin et al., 2012). Introns and their splicing add complexity to eukaryotic gene regulation: introns often contain cis-regulatory information that affects initiation, elongation, and/or termination steps of transcription (Dwyer et al., 2021; Gehring and Roignant, 2021). Alternative splicing can generate multiple protein isoforms from a single gene, which may differ in activity, function, and localization, and is often regulated in a cell-type-specific manner (Rogalska et al., 2023). Altered splicing patterns are also often associated with various human diseases, such as cancer (Bradley and Anczukow, 2023; Tao et al., 2024) and neurodegenerative diseases (Li et al., 2021).

Given the importance of splicing and its alteration in physiology and pathology, much effort has been devoted to analyzing splicing status. Various methodologies, from *in vitro* biochemical assays to RNA sequencing, have been employed to study splicing (Movassat et al., 2014). Short-and long-read RNA sequencing has been increasingly used to infer splicing efficiency and status, such as intron retention, exon skipping, alternative splicing, and mutually exclusive exons (Jiang et al., 2023; Prevorovsky et al., 2016; Sousa-Luis and Carmo-Fonseca, 2026; Wang and Rio, 2018; Wang et al., 2024).

The majority of approaches developed to analyze splicing events, such as rMATS (Wang et al., 2024), typically rely on predetermined structures of splicing events (e.g., use of alternative splicing sites, mutually exclusive exons, intron retention) based on genome annotation. That is, these computational programs include RNA-seq reads in their analyses only if the reads match predetermined, expected sequences of the splicing products. It has been recognized that these programs exclude a considerable number of RNA-seq reads from analysis, even though these excluded reads may contain biologically meaningful RNA species. Recognizing this limitation, other computational programs, such as Leafcutter (Li et al., 2018) and MAJIQ (Vaquero-Garcia et al., 2023), have been developed that do not rely solely on predetermining expected RNA sequences based on genome annotation. These programs have been used to detect RNA species that were not detectable by other approaches (Brown et al., 2022; Ma et al., 2022; Zeng et al., 2025). However, these programs focus on junction reads, in which two sequences are connected in linear order, thereby excluding non-canonical splicing events such as trans-splicing and back-splicing. Trans-splicing and back-splicing produce so-called chimeric reads, in which the two parts of RNA cannot be traced back to the genome sequence in a linear manner, and are discarded by most computational analyses. Detection of splicing events that yield such chimeric reads (e.g. back-splicing) typically requires a dedicated program (Vromman et al., 2023), making it difficult to detect their presence unless suspected already.

The need to detect non-canonical splicing events is especially acute under conditions that abundantly generate unexpected RNA species. We recently encountered this problem while analyzing the splicing of gigantic genes, which contain megabase-scale introns (Fingerhut et al., 2024; Fingerhut et al., 2019). Intron gigantism is observed across species, including mammals with ∼50 genes exceeding 1Mb due to gigantic introns (McCoy and Fire, 2020; Pozzoli et al., 2003; Pozzoli et al., 2002; Scherer, 2011). When the splicing of these gigantic introns was perturbed, a large amount of non-canonical RNA species were produced (Fontan et al., 2026). Capturing such events requires an approach that makes no prior assumptions about the RNA products that splicing should generate. To address the unmet need to detect and analyze non-canonical splicing events, we developed a new computational program, OmniSplice. Instead of relying on predefined splice-junction structures, OmniSplice first identifies RNA-seq reads that overlap an exon end (5’ or 3’), followed by the analysis of adjoining sequences. This approach allowed us to avoid discarding RNA-seq reads that do not match the anticipated RNA sequences. After recovering all RNA-seq reads that overlap with an exon end, the adjoining sequence is analyzed, allowing us to detect unanticipated RNA species in addition to expected RNA species. By applying OmniSplice to publicly available datasets [our previous datasets from *Drosophila* testes mutants for splicing factors (Fingerhut et al., 2024) and TDP-43 mutant mice that exhibit splicing phenotypes (Necarsulmer et al., 2023)], we detected splicing events and defects that had previously gone undetected, including constitutive back-splicing events that occur in wild-type conditions. Together, our results reveal that RNA-seq datasets may contain a substantial reservoir of overlooked splicing information. OmniSplice offers an approach to analyze splicing events without discarding non-canonical splicing products.

## Results

### Design of OmniSplice

Our motivation to develop OmniSplice stemmed from our previous studies on the expression of genes with ‘intron gigantism’ (Fingerhut et al., 2024; Fingerhut et al., 2019; Fontan et al., 2026). Intron gigantism, observed across metazoan genomes including mammals, is characterized by introns that can exceed hundreds of kilobases to megabases (Pozzoli et al., 2002; Scherer, 2011). Previously, we found that depletion of splicing factors (*U2af38* and *SRPK*) in the *Drosophila* testis increased defective splicing, which worsened with increasing intron size (Fingerhut et al., 2024). However, we suspected that the extent of splicing defects may not be fully captured by our previous analysis, because mis-splicing of gigantic introns may involve unexpected junctions that are missed by conventional splicing analysis programs. To detect unexpected splicing products, we designed OmniSplice to report and classify all RNA-seq reads that overlap the ends of exons. This method enabled us to detect previously missed splicing events.

OmniSplice utilizes two input files: the RNA-seq reads aligned to the reference genome as an indexed BAM file and the genome annotation as a GTF file. To capture defective splicing events, it is desirable to use a total RNA-seq dataset, although a polyA-selected RNA-seq dataset may also yield some insights. From the GTF file, OmniSplice extracts all introns and uses them to build an interval tree (a data structure optimized to recover overlapping reads, Fig. 1A). Then, OmniSplice reads the BAM file, recovering RNA-seq reads that overlap exon ends (5’ or 3’) by querying the interval tree, and filtering out any RNA-seq reads that do not overlap the exon ends and those that fail quality control (secondary alignment) (Fig. 1A). If the RNA-seq reads are stranded, OmniSplice categorizes ‘wrong strand’ in a separate category: we normally do not analyze this category further, but the counts are recorded by OmniSplice, and thus can be further analyzed if deemed necessary (e.g. if the user suspects cryptic transcription in the wrong direction).

**Fig 1.**
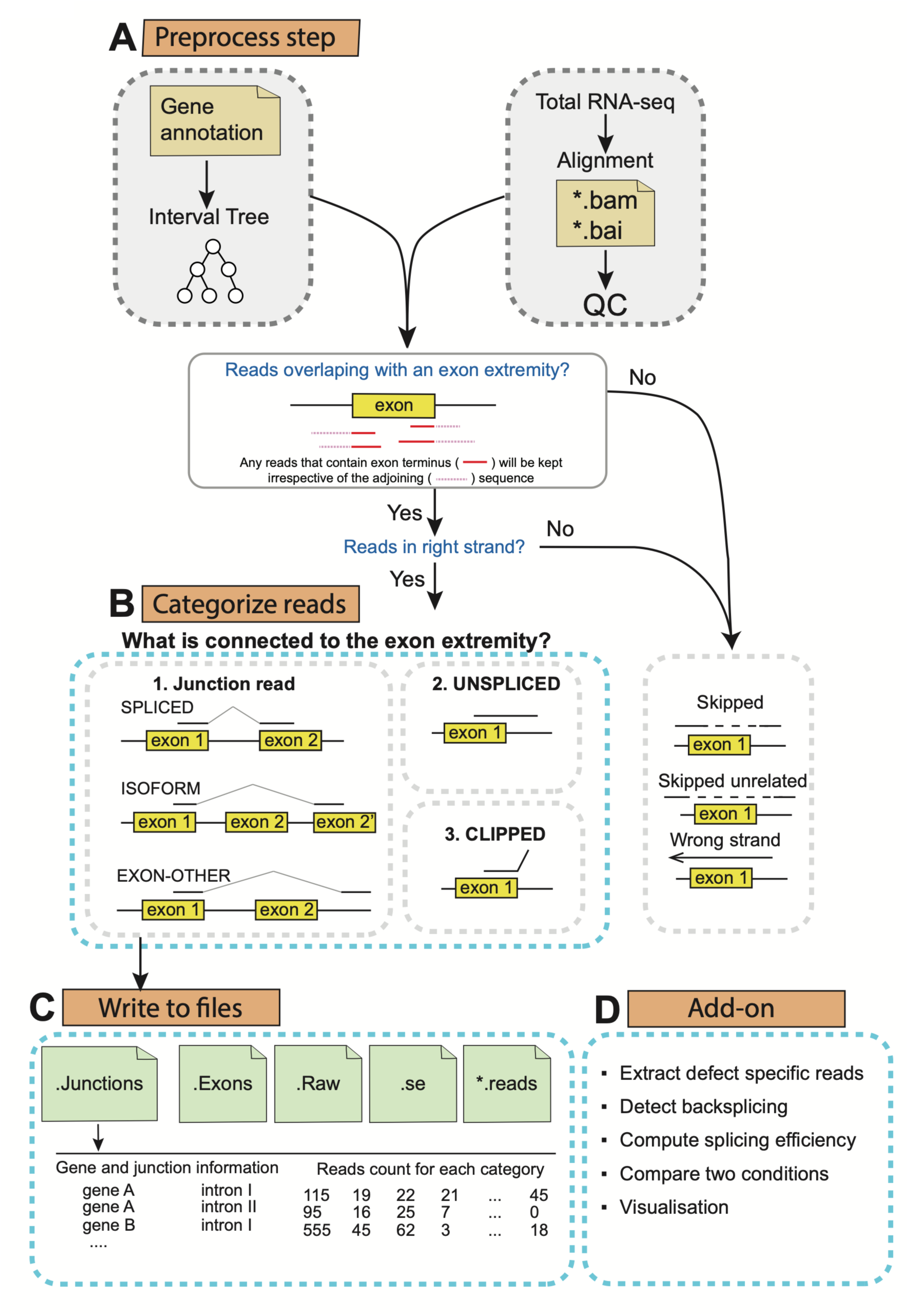
OmniSplice workflow. OmniSplice requires two inputs: 1) a sorted, indexed BAM file from a total-RNA-seq experiment. 2) a gene annotation as a GTF file. OmniSplice applies the following procedure for each read. First OmniSplice filters out reads that fail quality control. By default, OmniSplice QC filters out reads with mapq < 13 and uses the SAM flag to filter out reads marked as1) supplementary alignment, 2) PCR duplicate, 3) not primary alignment, or 4) failing the platform quality check. Then OmniSplice identifies reads that overlap any exon end (5’ side or 3’ side of an exon) by using an interval tree built from the gene annotation. In the case of stranded RNA-seq, OmniSplice discards reads from the wrong (anti-sense) strand. For every exon, OmniSplice categorizes the read based on the sequence adjoining the exon. The counts of RNA-seq reads in each category for each exon are written to output files. Optionally (depending on the user’s interest), reads from a specific category can be written to a separate file for further inspection. The information in those files can be used for downstream analyses. We have also generated pre-made, add-on analysis tools that allow detection of back-splicing, computation of splicing efficiency, and statistical comparison between two conditions.

The RNA-seq reads that overlap exon ends and passed the quality control will be further analyzed based on the sequence adjacent to the exon ends (referred to as ‘adjoining sequence’) (Fig. 1B). Importantly, any RNA-seq reads that overlap exon ends are included at this step, allowing the recovery of RNA-seq reads that may be produced by unexpected and/or erroneous splicing events. RNA-seq reads are generally categorized as follows:

1. Junction read: an exon end is connected to a sequence in a manner that allows linear (non-chimeric) alignment. Junction reads are further categorized as follows.

i. SPLICED: an exon end is connected to another exon in a manner that indicates correct splicing based on genome annotation.
ii. ISOFORM: an exon end is connected to another exon, compatible with a known alternative isoform.
iii. EXON-OTHER: an exon end is connected to a sequence that is not expected from the genome annotation. This category may include previously unreported isoforms or erroneous splicing (such as an exon being connected to the middle of another exon or an intron).
2. UNSPLICED: the RNA sequence matches that of the genome sequence, indicating no splicing occurred.
3. EXON-CLIPPED: The RNA-seq read was aligned up to the exon end, but the adjoining sequence is marked as ‘clipped’ by the aligner. The aligner program clips the reads when the adjoining sequence is not found anywhere in a manner that allows linear (non-chimeric) alignment. The adjoining sequence may map to another chromosome or the same chromosome (see below).

Most splicing analysis programs typically discard RNA-seq reads in categories of EXON-OTHER and EXON-CLIPPED, assuming they are likely due to experimental artifacts. However, OmniSplice is designed to retain these reads as well, allowing subsequent deeper analysis as necessary. Note that the categories used by OmniSplice will be indicated by capital letters (e.g. EXON-OTHER, SPLICED) throughout the manuscript to distinguish them from general descriptions of splicing events.

After analyzing RNA-Seq reads using these criteria, OmniSplice outputs three main files (Fig.1C):

1. .exons: details the read count at each exon end, categorizing them based on adjoining sequence (Fig.1B) as described above.
2. .junctions: details the read counts of splicing events. The method of analyzing the exon ends is the same as above (.exons), but the above method counts a single RNA-Seq read twice if two exon ends are joined. The.junctions file is the result of removing such double counts from the.exons file (see Fig. S1 for the definition of.exons vs.junction).
3. .raw: unprocessed raw output, counting individual events, and giving genomic position of spliced RNA-Seq reads. This file is useful for subsequent analyses, when the exact splicing events need to be analyzed.

Based on these outputs, further analysis can be carried out. For example, splicing efficiency can be calculated based on spliced reads/(spliced reads + unspliced reads). OmniSplice provides a default program module to compute the splicing efficiency in this manner, with the results provided as an.se file. Alternative calculation methods can be applied flexibly based on users’ needs: for example, the frequency of unexpected splicing events can be calculated using [1 – (SPLICED and UNSPLICED)/total reads]. Optionally, OmniSplice can output files with reads from specific categories for downstream analyses (*reads file). OmniSplice also provides pre-made scripts (Fig. 1D) for result visualization, back-splicing detection (see below), and statistical analyses.

Because OmniSplice simply categorizes all exon-end-containing reads by the connected sequence and counts each event, the output data file (provided as tabulated read counts in the.tsv format) is amenable to downstream analyses at the users’ discretion. For example, we further analyzed RNA-seq reads in the EXON-CLIPPED category in our recent study using sterile hybrids between *D. simulans* and *D. mauritiana* (Fontan et al., 2026). Although EXON-CLIPPED reads are often considered the result of technical artifacts (e.g. ligation of two unrelated RNA molecules during library prep), our study found EXON-CLIPPED reads that represent biological ‘back-splicing’ events (the end of an exon is connected to the beginning of an earlier exon), highlighting the usefulness of the OmniSplice approach.

### Comparison of OmniSplice with rMATS and MAJIQ

We previously generated stranded total RNA-seq datasets following RNAi-mediated depletion of splicing factors (*SRPK, U2af38*) in the male germline of *D. melanogaster, and* found that depletion of these genes led to aberrant splicing of many genes, particularly those with gigantic introns (Fingerhut et al., 2024). A handful of genes in the genome from flies to humans contain genes with gigantic introns that can exceed hundreds of kilobases to even megabases (Fingerhut et al., 2019; Scherer, 2011). In *Drosophila*, such gigantic genes are mostly found on the Y chromosome and expressed during spermatogenesis. While we found some aberrant splicing patterns (Fingerhut et al., 2024), we suspected that many more aberrant/erroneous splicing events may be missed, considering the challenging nature of splicing across gigantic introns (Fingerhut et al., 2024; Fingerhut et al., 2019).

Using this dataset, we compared OmniSplice with rMATS (rMATS-turbo)(Wang et al., 2024) and MAJIQ(Vaquero-Garcia et al., 2023), widely utilized programs that analyze splicing status. rMATS detects changes in splicing patterns between two conditions and identifies canonical alternative splicing event (i.e. annotated splicing events, including alternative splice site usage, skipped exon, intron retention). MAJIQ is also designed to detect changes in the splicing patterns without solely relying on previously annotated splicing junctions. However, neither rMATS or MAJIQ is designed to detect chimeric RNA species that join two pieces of DNA in a non-linear manner (e.g. back-splicing and certain trans-splicing). On the other hand, OmniSplice is designed to recover all RNA-seq reads that contain exon ends, identifying unexpected junction reads.

When the RNA-seq data of *SRPK^RNAi^* and *U2af38^RNAi^*were analyzed, OmniSplice recovered more events than rMATS and MAJIQ (Fig. 2A). RNA-seq reads that were uniquely identified by OmniSplice spanned all categories of OmniSplice classification (Fig. 2B, C). The UNSPLICED category is expected to be unique to OmniSplice, because rMATS and MAJIQ do not detect this category by design: they only detect ‘intron retention’, which is computed by a different definition (i.e. coverage of the intronic sequence). The increase in the UNSPLICED category detected by OmniSplice shows that *SRPK^RNAi^* and *U2af38^RNAi^*conditions have reduced splicing efficiency. Although this is not surprising for the perturbation of splicing factors (Allemand et al., 2001; Graveley, 2000; Shao et al., 2014; Zamore and Green, 1989), it reveals the utility of OmniSplice in automatically detecting decreased splicing efficiency, without using a separate splicing efficiency program. Most notable category that is uniquely identified by OmniSplice is EXON-CLIPPED (Fig. 2B, C). The majority of splicing analysis programs are designed to ignore clipped part of RNA-seq reads, thereby entirely excluding junctions that belong to the EXON-CLIPPED category, whereas OmniSplice reports them, contributing to the detection of RNA-seq reads that are missed by other programs (see below for details). Thus, OmniSplice offers a complementary approach to other existing programs to analyze splicing.

**Fig 2.**
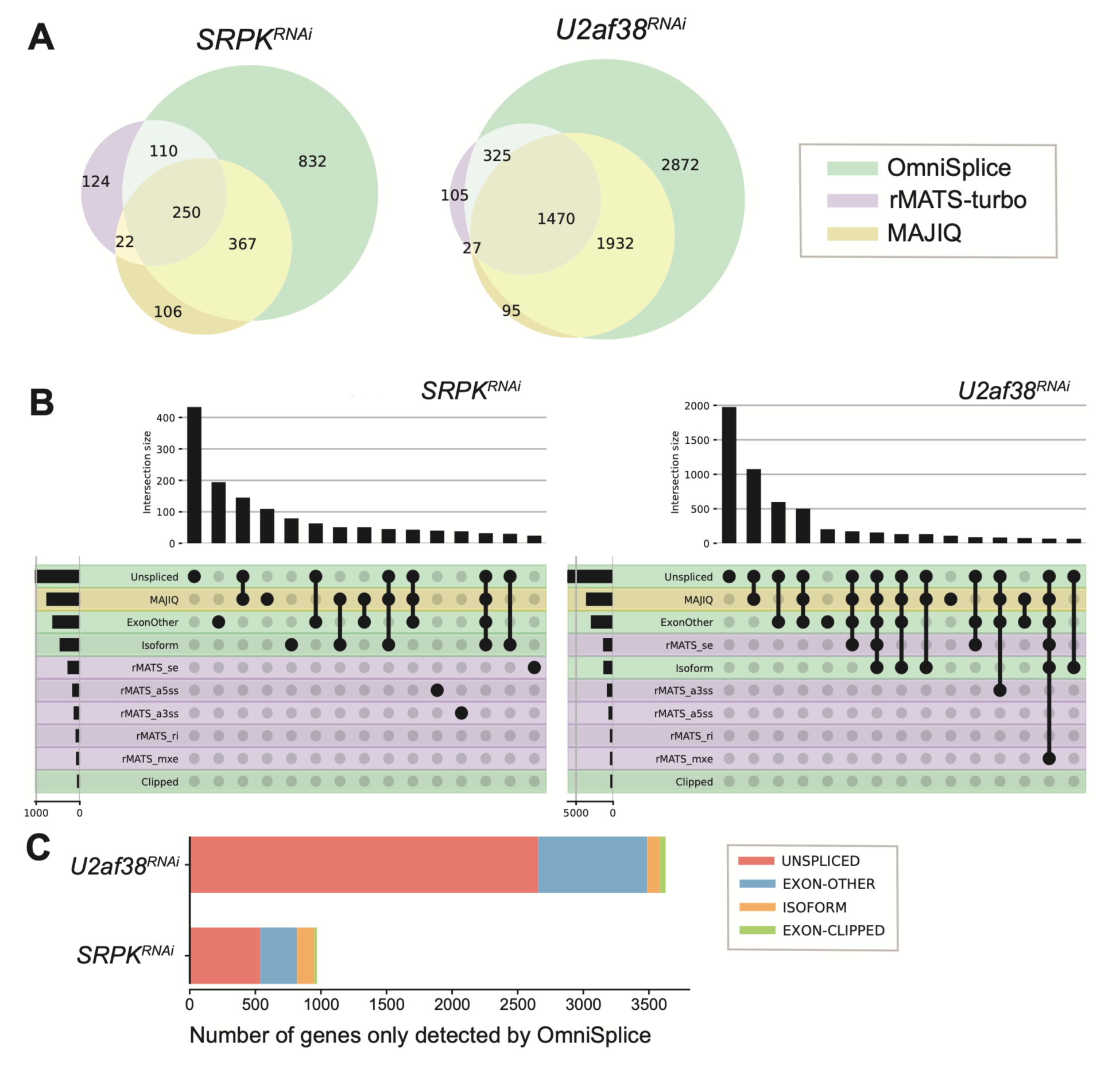
Comparison of OmniSplice, rMATS and MAJIQ in recovering altered splicing in the datasets of *Drosophila* splicing mutants. A) Venn diagram of genes recovered by OmniSplice (green), rMATS (purple) and MAJIQ (yellow) as exhibiting significantly altered splicing status (adjusted p-value <0.01 and |delta-psi| > 1), comparing control and *SRPK^RNAi^*(left) or *U2af38^RNAi^* (right). B) UpSet plots (Lex and Gehlenborg, 2014) to compare the distribution and overlap between the rMATS, MAJIQ and OmniSplice categories for recovered genes in *SRPK^RNAi^* (left) and *U2af38^RNAi^* (right). A gene may be counted in more than one category if it contains junctions assigned to different categories. The x-axis is sorted by intersection size and only the top 12 intersections are shown. Row color indicates the origin of analysis tools for each category: rMATS (purple), MAJIQ (yellow) or OmniSplice (green). E) The categories of genes recovered only by OmniSplice (corresponding to the green area in Fig. 2A). Note that a gene may be presented in multiple categories, when they have multiple introns exhibiting different categories of defects.

### OmniSplice detects back-splicing events

We further analyzed the RNA-seq reads that were uniquely identified by OmniSplice in *SRPK^RNAi^* and *U2af38^RNAi^*conditions, particularly focusing on the reads in the EXON-CLIPPED and EXON-OTHER categories (Fig 2C). We plotted genes based on the strength of change between RNAi and control (using delta-psi on X-axis, see methods) and the associated adjusted p-values (Y-axis), visualizing as volcano plots (Fig 3A). This analysis revealed considerable changes in the splicing patterns in *U2af38^RNAi^*and *SRPK^RNAi^* conditions compared to controls (Fig 3A). In both conditions, the Y-linked genes were markedly impacted, as indicated by our previous study (Fig 3A)(Fingerhut et al., 2024).

**Fig 3.**
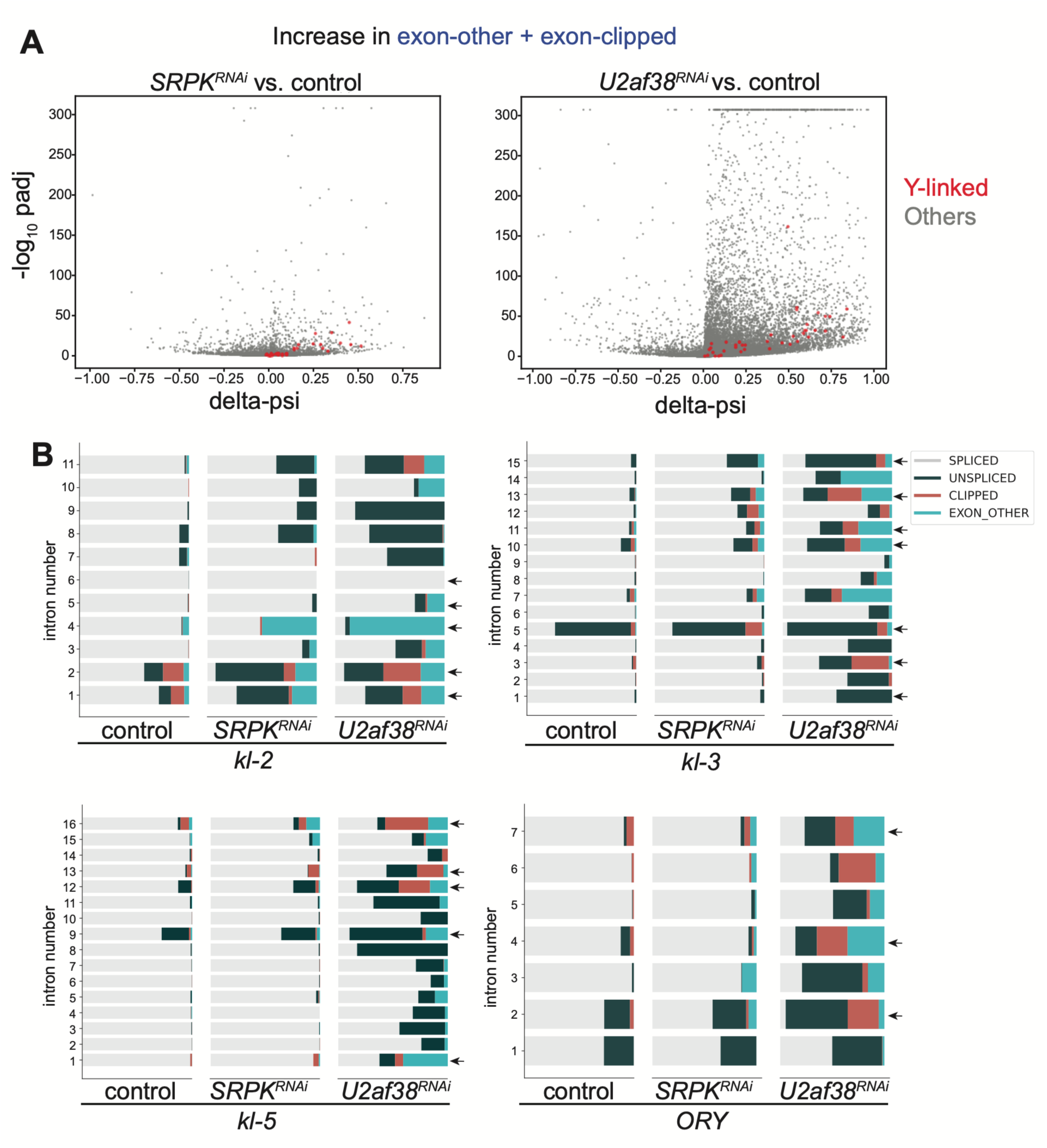
OmniSplice reveals widespread splicing defects in *Drosophila* splicing factor mutants. A) Volcano-style plots for *SRPK^RNAi^* (left) and *U2af38^RNAi^* (right), showing the delta-psi (X-axis) and the adjusted p-values using Benjamini-Hochberg method (Y-axis). The p-value and ratio values were computed using OmniSplice ‘*compare_conditions’* module (see materials and methods). For this analysis, the ‘*compare_conditions’* module was used to compare the ratio of “SPLICED / (EXON-OTHER + EXON-CLIPPED)” between the genotypes for every junction. B) Bar plots showing the splicing status of Y-linked gigantic genes in control, *SRPK^RNAi^*and *U2af38^RNAi^*. Each row represents individual introns, where the proportion of reads from the different categories is shown: SPLICED (light grey), UNSPLICED (dark green), EXON-CLIPPED (orange), EXON-OTHER (blue). Arrows indicate gap-containing introns.

Y-linked genes are characterized by their gigantic introns, some of which exceed megabases. We have shown that gigantic introns pose a challenge in splicing, making them sensitive to perturbations (Fingerhut et al., 2024; Fontan et al., 2026). We analyzed individual Y-linked genes at each exon-intron junction using OmniSplice (Fig 3B, Fig S2). This analysis revealed splicing defects of the Y-linked genes in the *U2af38^RNAi^*and *SRPK^RNAi^* condition, beyond our previous report (Fingerhut et al., 2024). The increase in the UNSPLICED, EXON-OTHER, and EXON-CLIPPED categories was clearly identified in *U2af38^RNAi^* (Fig 3B, Fig S2). *SRPK^RNAi^* also exhibited widespread splicing defects, albeit to a lesser extent compared to *U2af38^RNAi^*(Fig 3B, Fig S2). The aberrant splicing (belonging to the EXON-OTHER and EXON-CLIPPED categories) increased considerably at the gigantic introns containing genome assembly gaps in the *SRPK^RNAi^* and *U2af38^RNAi^*conditions (Fig 3B, Fig S2, gigantic introns are marked by arrows), reinforcing the conclusion that the splicing of gigantic introns is particularly challenging, as suggested previously (Fingerhut et al., 2024; Fontan et al., 2026).

In our previous study that analyzed hybrids between *D. simulans* and *D. mauritiana*, we observed a considerable increase in EXON-CLIPPED reads using an early version of OmniSplice (Fontan et al., 2026). Many of the EXON-CLIPPED reads were due to back-splicing, where the end of an exon (typically right before a gigantic intron) was joined to the beginning of an earlier exon (Fontan et al., 2026). Thus, we investigated whether the EXON-CLIPPED reads that were increased in the Y-linked gigantic genes in the *SRPK^RNAi^*and *U2af38^RNAi^* conditions were also due to back-splicing. Indeed, we found RNA-seq reads consistent with back-splicing, and after filtering out events with less than 5 reads in each replicate, we identified a clear signature of back-splicing in 6, 9 and 47 junctions in the control, *SRPK^RNAi^*and *U2af38^RNAi^* conditions, respectively (Table S1). Among these, back-splicing of *kl-2* intron 2 was confirmed by RT-PCR, followed by direct sequencing of the PCR products (Fig S3).

OmniSplice also detected back-splicing events outside the Y-linked gigantic genes: by analyzing the EXON-CLIPPED reads, we found constitutive back-splicing events in autosomal genes, including *CAP, mbl, stw* and *nrv2* (Table S1). The back-splicing events in these genes were observed at introns that are relatively large (4.4 – 75 kb) but not gigantic (Table S1, Fig. S4). Interestingly, these constitutive back-splicing events were more frequently observed in control than in splicing mutants (Table S1, Fig. S4A, B), suggesting that these back-splicing events may be biologically relevant and positively regulated by splicing factors. These constitutive back-splicing events observed in the wild type may represent regulatory circular RNAs (Chen, 2020). Back-splicing of *CAP* and *stw* genes was also confirmed by RT-PCR, followed by the sequencing of the PCR products (Fig. S4).

Importantly, the constitutive back-splicing observed in wild type conditions would have been missed by rMATS and MAJIQ, revealing the utility of OmniSplice program. Detection of back-splicing typically requires a dedicated program (Vromman et al., 2023). Accordingly, back-splicing normally goes unnoticed unless suspected already. In contrast, because OmniSplice recovers all RNA-seq reads that overlap with the exon ends, back-splicing events are automatically included in the EXON-CLIPPED category. Thus, OmniSplice serves as an easy entry point by flagging ‘unusual’ RNA species, prompting further analysis.

### OmniSplice detects trans-splicing events in the *Drosophila* testis

In addition to EXON-CLIPPED reads (including back-splicing events as described above), we noted a considerable increase in RNA-seq reads in the EXON-OTHER category, particularly in the *U2af38^RNAi^* condition (Fig. 2C, Fig 3B, Fig S2-4), prompting us to further investigate the nature of these RNA-seq reads in more detail. We found that the increase in the EXON-OTHER category was not limited to the Y-linked gigantic genes, and we observed global increases in this type of RNA-seq reads in *SRPK^RNAi^* and *U2af38^RNAi^* conditions (Fig 4A, B). Focusing on the EXON-OTHER reads that were significantly increased in the RNAi conditions compared to the control (orange dots in Fig 4B), we further subcategorized the RNA-seq reads in the EXON-OTHER category (Fig 4C). Surprisingly, we found that some of the EXON-OTHER reads represented splicing events between two genes (Fig 4C). This indicates trans-splicing between two RNA molecules. Alternatively, a large region of DNA may be transcribed together as a single transcript (e.g. read-through transcription due to failure in transcription termination), which is then spliced to yield ‘chimeric RNA’ between two genes. Considering that such RNA-seq reads indicative of trans-splicing increased specifically in *SRPK^RNAi^* and *U2af38^RNAi^* conditions (Table S2), it is unlikely that trans-splicing reads are due to a mere artifact of library preparation. Indeed, the presence of trans-spliced molecules was confirmed by RT-PCR, followed by sequencing (Fig S5).

**Fig 4.**
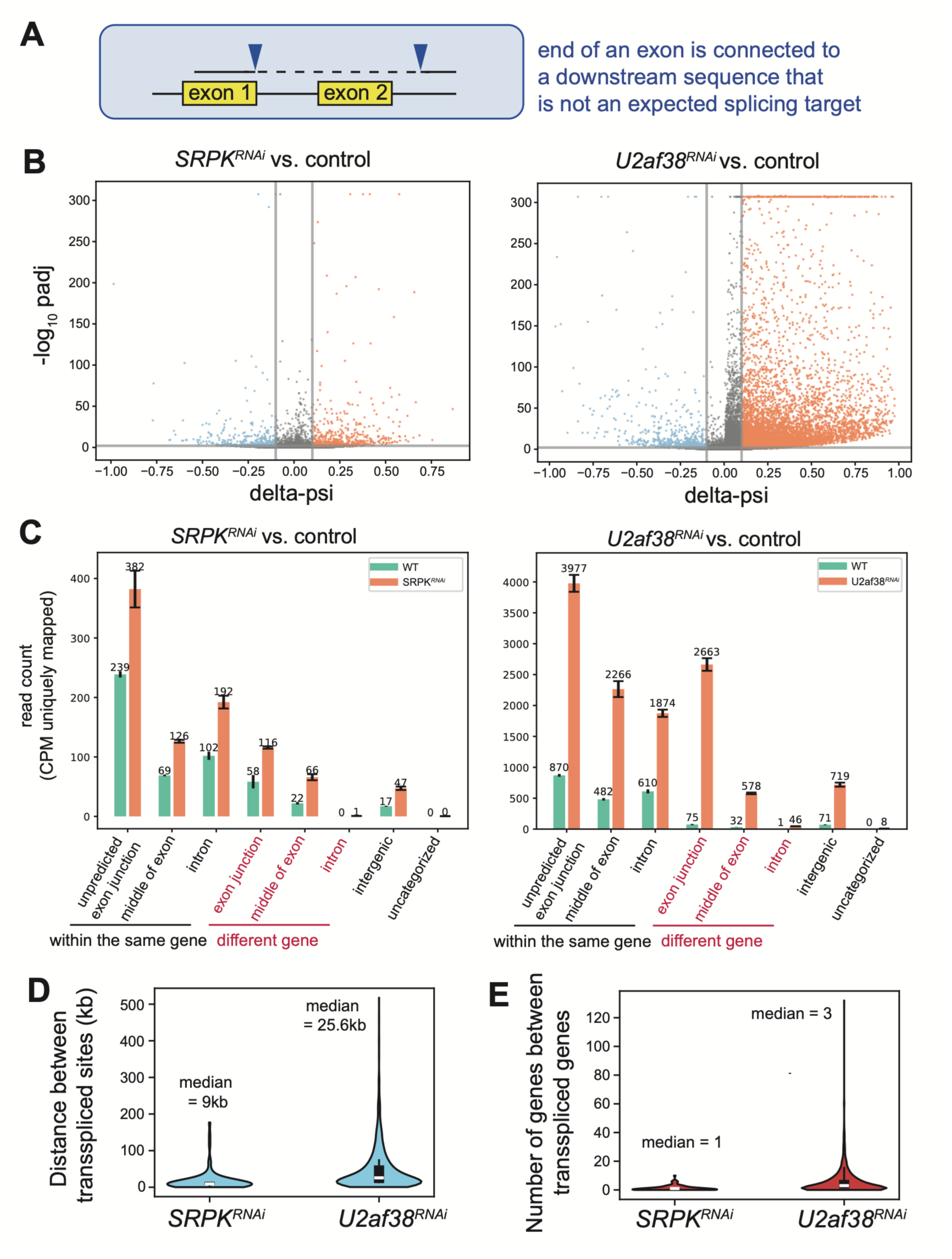
Analysis of ‘EXON-OTHER’ reads by OmniSplice reveals widespread defects and potential trans-splicing events in *Drosophila* splicing factor mutants. A) Schematic example of an EXON-OTHER read. B) Volcano-style plots for the EXON-OTHER category in *SRPK^RNAi^* (left) and *U2af38^RNAi^* (right), showing the delta-psi (X-axis) and the adjusted p-value (Benjamini-Hochberg procedure) (Y-axis). The p-value and ratio values were computed using the OmniSplice “*compare_conditions*” module, comparing the ratio “SPLICED / (SPLICED + EXON-OTHER)” between conditions for every junction. Junctions significantly enriched in the RNAi and control (adjusted p-value < 0.01 and |delta-psi| > 0.1) are highlighted in orange and blue, respectively. C) Normalized read counts (per million uniquely mapped reads) of the EXON-OTHER category (divided into 8 subcategories) in *SRPK^RNAi^* (left) and *U2af38^RNAi^*(right). Each EXON-OTHER read found to be significantly increased in the RNAi condition (orange dot panel B) was assigned to one of eight subcategories according to where the adjoining sequence mapped. Orange bars represent RNA-seq reads increased in the RNAi condition, while green bars indicate the corresponding RNA-seq reads in the control. (D) Distance between two genes that are potentially ‘trans-spliced’ in the category of ‘different gene – exon junction’ category in (C). (E) The number of genes between the two genes that are potentially ‘trans-spliced’ in the category of ‘different gene – exon junction’ category in (C).

We further analyzed the relationship between two genes that are spliced together. The average distance between two genes was 26kb in *U2af38^RNAi^*, and 10.5 kb in *SRPK^RNAi^*, with some cases reaching >100kb (Fig. 4D). Also, on average, there were 3 genes in *U2af38^RNAi^*, and 1 gene in *SRPK^RNAi^*(up to 10s of genes in some cases) between two genes that produced chimeric reads (Fig. 4E). By examining individual cases of ‘two-gene chimera’, we found that the intervening genes were often not transcribed at all (examples from top hits are shown Fig S6). Based on these observations, we favor trans-splicing as the cause of producing EXON-OTHER reads that connect two genes, although other possibilities cannot be entirely excluded.

### Comparison of OmniSplice with rMATS and MAJIQ in the analysis of the TDP-43 mutant mice

While the above analysis revealed the utility of OmniSplice in detecting non-canonical RNA species that escape the detection of conventional splicing analysis programs, *U2af38^RNAi^* and *SRPK^RNAi^*conditions are extreme examples of splicing perturbation. Thus, to further test the utility of OmniSplice in detecting more subtle changes, we analyzed publicly available RNA-seq datasets that compared gene expression between wild-type and TDP-43 K145Q mutant mice (TDP-43 KQ, hereafter) in the cortex as well as the hippocampus (Necarsulmer et al., 2023). TDP-43 KQ mutant condition is expected to exhibit much more subtle defects in splicing, compared to knockdown of splicing factors (SPRK and U2af38), allowing us to test the performance of OmniSplice.

TDP-43 is a multifunctional RNA-binding protein and is known to play important roles in splicing regulation (Tollervey et al., 2011). Dysregulation or mutation of TDP-43 has been identified in numerous patients with neurodegenerative diseases such as Amyotrophic Lateral Sclerosis (ALS) and Frontotemporal Dementia (FTD)(Cairns et al., 2007; Hogan et al., 2016; Neumann et al., 2007; Neumann et al., 2006; Prater et al., 2022). The TDP-43 KQ mutation has been shown to mimic acetylation at Lysine 145, a modification often found in pathological TDP-43 protein inclusions associated with multiple neurodegenerative diseases (Cohen et al., 2015; Wang et al., 2017). The TDP-43 KQ mouse model recapitulated several phenotypes associated with human neurogenerative disease, including TDP-43 cytoplasmic mislocalization, splicing dysfunction, and cognitive impairment (Necarsulmer et al., 2023).

Using the wild-type and TDP-43 KQ datasets (stranded total RNA-seq data from the cortex and hippocampus of control and TDP-43 KQ female mice in triplicate) (Necarsulmer et al., 2023), we compared OmniSplice with those from rMATS and MAJIQ. First, we compared the number of genes whose splicing status was found to be significantly different (q-value < 0.01 and |delta-psi| > 0.1) between control and TDP-43 KQ mutant by either OmniSplice, rMATS or MAJIQ. In both cortex and hippocampus samples, OmniSplice identified more genes than rMATS or MAJIQ (Fig 5A). The RNA-seq reads that were uniquely identified by OmniSplice spanned all categories of OmniSplice classification (Fig. 5B, C). The genes identified by the three programs exhibited less overlap, compared to the analysis of *U2af38^RNAi^* and *SRPK^RNAi^* conditions. Identification of overlapping but distinct sets of differentially-spliced genes by these three programs likely reflects the different approaches these programs use to define altered splicing. A similar observation was made in studies comparing multiple splicing analysis programs (Draper et al., 2025; Jiang et al., 2023).

**Fig 5.**
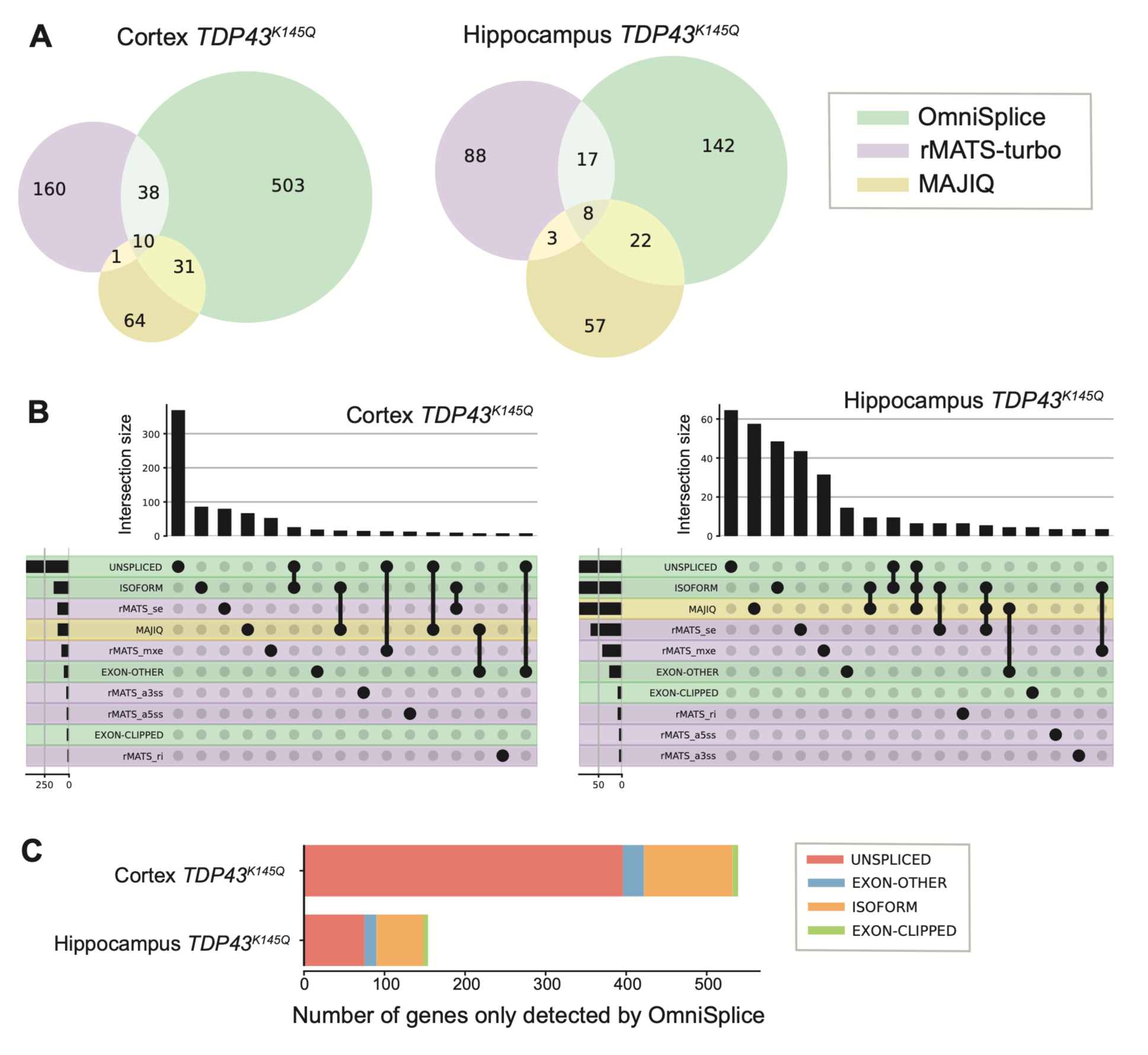
Comparison of OmniSplice, rMATS and MAJIQ in recovering altered splicing in the datasets of mouse TDP43^K145Q^ mutant. A) Venn diagram of genes recovered by OmniSplice (green), rMATS (purple) and MAJIQ (yellow) as exhibiting significantly altered splicing status (adjusted p-value <0.01 and |delta-psi| > 0.1), comparing control and TDP43^K145Q^ mutant in the cortex (left) and Hippocampus (right). B) UpSet plots (Lex and Gehlenborg, 2014) to compare the distribution and overlap between the rMATS, MAJIQ and OmniSplice categories for recovered genes in the cortex (left) and the hippocampus (right). A gene may be counted in more than one category if it contains junctions assigned to different categories. The X-axis is sorted by intersection size and only the top 12 intersections are shown. Row color indicates the origin of analysis tools for each category: rMATS (purple), MAJIQ (yellow) or OmniSplice (green). E) The categories of genes recovered only by OmniSplice (corresponding to the green area in Fig. 5A). Note that a gene may be presented in multiple categories, when they have multiple introns exhibiting different categories of defects.

### OmniSplice identifies cryptic/unannotated exons in the TDP-43 mouse model

It is known that TDP-43 mutation/pathology increases cryptic exons, intronic sequences that are erroneously spliced into RNA products (Brown et al., 2022; Klim et al., 2019; Ling et al., 2015; Ma et al., 2022; Melamed et al., 2019). Because OmniSplice is designed to detect non-canonical splicing sites that involve a sequence that is not annotated as an exon, we tested whether OmniSplice could recover cryptic exons in the TDP-43 KQ mutant.

By applying OmniSplice to the TDP-43 KQ mutant datasets (Necarsulmer et al., 2023), we found an increase in the EXON-OTHER category in the TDP-43 KQ mutant (Fig 6A), whereas we did not find a marked increase in the EXON-CLIPPED categories (Fig S7). The UNSPLICED category also increased (Fig S7), suggesting that TDP-43 KQ mutant exhibits reduced splicing efficiency. We further investigated the nature of RNA-seq reads in the category of EXON-OTHER in the TDP-43 KQ mutant. We found that the majority of the EXON-OTHER category reads that increased in the TDP-43 KQ mutant represent the splicing events from the end of an exon to either the middle of another exon or intron of the same gene (Fig 6B). It is interesting to note that there were barely any detectable EXON-OTHER category reads that indicate trans-splicing, as was observed upon depletion of splicing factors in the *Drosophila* testis (Fig 4C).

**Fig 6.**
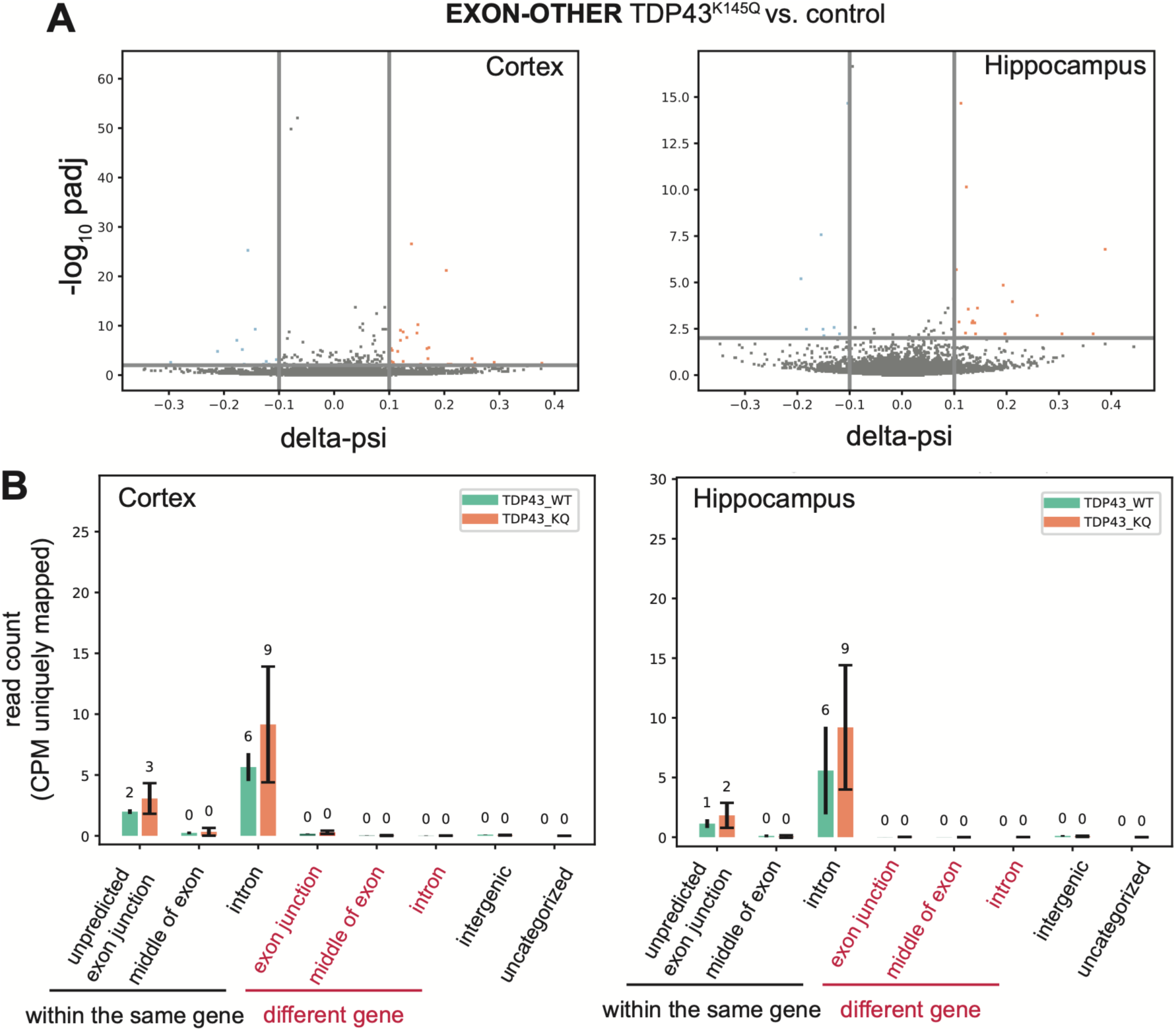
Analysis of the EXON-OTHER category reads in the TDP-43 KQ mutant datasets by OmniSplice reveals intragenic splicing defects. A) Volcano-style plot for the EXON-OTHER category in the cortex (left) and the hippocampus (right) in TDP-43 KQ conditions, showing the delta-psi (X-axis) and the adjusted p-value (Benjamini-Hochberg procedure) (Y-axis). The p-value and ratio values were computed using OmniSplice “*compare_conditions*” module, comparing the ratio “SPLICED / (SPLICED + EXON-OTHER)” between conditions for every junction. The delta-psi provides the effect size for ratio changes (see Material and Methods). Junctions significantly enriched in the TDP-43 KQ and control (adjusted p-value < 0.01 and | delta-psi | > 0.1) are highlighted in orange and blue, respectively. B) Normalized read counts (per million uniquely mapped reads) of the EXON-OTHER category (divided into 8 subcategories) in the cortex (left) and the hippocampus (right). Each EXON-OTHER read found increased in the mutant condition (orange dot panel B) was assigned to one of eight subcategories according to where the adjoining sequence mapped. Orange bars represent RNA-seq reads increased in the TDP-43 KQ mutant condition, while green bars indicate the corresponding RNA-seq reads in the control.

By further examining these non-canonical splicing junctions found in TDP-43 KQ mutants, we observed many splicing events that join the end of an exon to somewhere in the middle of an intron, as well as those that join somewhere in the middle of an intron to the beginning of an exon. Combining these two events often led to increased read coverage between two non-canonical splicing sites, indicating that a cryptic exon was indeed included in the mature RNA (Fig 7A, *Arhgap10* gene). We also observed cases where the cryptic exon appears to exist to some extent even in the control (Fig 7B, C, D), suggesting that those sequences have a tendency to be mis-spliced even in wild-type conditions, which is exacerbated in the TDP-43 KQ mutant (Fig 7B, C). The cryptic exons in genes *Golt1b* and *Sema4d* (Fig 7B, C) would cause frame-shift, if included. In contrast, in the case of the *Rims2* gene (Fig 7D), the inclusion of the ‘cryptic exon’ produces an in-frame ORF, suggesting that it might be an unannotated exon (constitutive or alternative) that is a part of the normal mRNA product. Finally, although the presence of UG repeats was reported to be present near the cryptic exons (Ling et al., 2015), cryptic exons found by OmniSplice did not always have UG repeats nearby (red arrows in Fig 7), implying that the mis-splicing induced by TDP-43 mutant may be more widespread than previously appreciated. Among these examples, cryptic exons for *Arhgap10, Sema4d* and *Rims2* were identified by MAJIQ as well, while that of *Golt1b* was only identified by OmniSplice.

**Fig 7.**
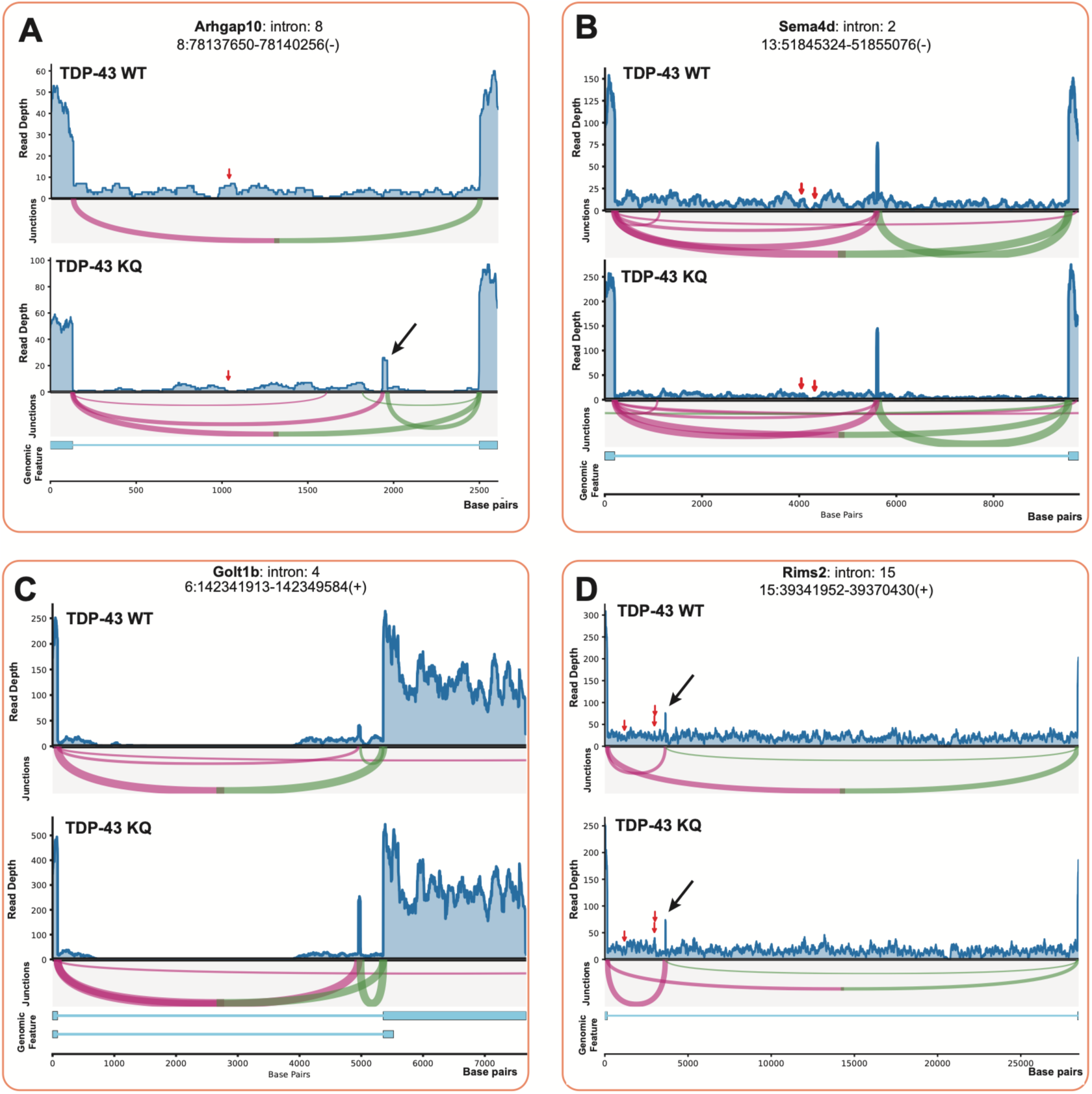
OmniSplice detects potential cryptic and unannotated exons in the TDP43^K145Q^ cortex dataset. Sashimi plots of potential cryptic exons for Arhgap10 (A), Sema4d (B), Golt1b (C), and Rims2 (D). The exon indicated by the black arrow in A (whose read increases specifically in the TDP43^K145Q^ mutant condition) introduces a stop codon, indicating this is an erroneously generated cryptic exon. The exon indicated by the black arrow in D, which is seen both in the control and the TDP43^K145Q^ mutant conditions, introduces an in-frame addition of 9 amino acids, implying that this may be an unannotated, correct exon. The Y-axis represents the raw read coverage. In the sashimi plot, the arc width and Y-position are proportional to the normalized read depth. The genomic-feature tracks represent annotated genomic features. Red arrow indicates the presence of a UG array (≥ 7 repeats).

## Discussion

Motivated by our need to detect splicing defects that produce RNA species that do not fall into predetermined categories of splicing events, we developed OmniSplice, which is designed to detect all RNA-seq reads that overlap exon ends, agnostic of ‘what sequence is adjoined to the end of an exon’. This approach enabled us to recover many RNA-seq reads that are typically discarded by other programs designed to analyze splicing events. The notable advantage of OmniSplice is the detection of chimeric reads, which are typically discarded from the analysis by many programs. OmniSplice allowed us to discover back-splicing events as well as potential trans-splicing events. Importantly, OmniSplice detected back-splicing events that are present in wild type conditions, which would have been missed by other programs that exclude non-linear alignments (chimeric reads that belong to the EXON-CLIPPED category of OmniSplice). Although the dedicated programs to detect back-splicing exist, analysis to search for back-splicing events are often not conducted unless already suspected. Thus, OmniSplice serve as a convenient ‘one-stop shop’ analysis tool to simultaneously examine various splicing status, such as splicing efficiency, alternative splicing and back-splicing.

OmniSplice was able to discover splicing changes in TDP-43 mutant mice that exhibit much more subtle splicing defects compared to *SPRK* and *U2af38* mutant conditions. In analyzing TDP-43 mutant mice dataset, OmniSplice performed similarly to rMATS and MAJIQ. Together, OmniSplice offers an orthogonal approach to analyzing splicing defects, particularly suitable for identifying non-canonical splicing events that create chimeric reads, which are ignored by most software designed to analyze alternative splicing. More importantly, our results suggest that RNA-seq datasets contain a substantial reservoir of overlooked splicing information, warranting more comprehensive approaches to analyze RNA-seq data for splicing events.

## Methods

### OmniSplice

GitHub: https://github.com/whitehead/OmniSplice release: “v0.5.0 April 2026, submission release”. Code to generate figures and analyses has been added to https://github.com/rLannes/OmniSplice_Analyses_Figures. OmniSplice is written in rust and depends on the following crate: bio 1.0, rust-htslib 0.45.0, CigarParser (https://github.com/rLannes/CigarParser), clap 4.5.16, strand_specifier (https://github.com/rLannes/BAMstrandSpecifier), regex, lazy_static, itertools, log “0.4”, flexi_logger “0.31”, thiserror, nalgebra, rayon 1.10:

### Other software used

Data analysis and plotting were performed with Python v3.12 and Rust code (rustc 1.95.0). For data manipulation, we used pandas v2.2.2, intervaltree v3.1.0, numpy v2.0.1, scipy v1.14.0. For visualization we used matplotlib v3.10.7. Several custom libraries were developed for this study: easyfasta v.1.1.64 (https://github.com/rLannes/easyfasta) was used to parse fasta file. gtf_pyparser v0.0.0 (https://github.com/rLannes/gtf_pyparser) was used to work with gtf. pycoverplot v0.1.0(https://github.com/rLannes/pycoverplot) was used to make RNA-seq reads coverage plot. Following two Rust backends, integrated with Python via maturin and PyO3, were used for performance-critical operations on BAM files: Rust_covpyo3 (https://github.com/rLannes/Rust_covpyo3) for recovering RNA-seq reads coverage from BAM files over a genomic interval, and Rust_get_junctions (https://github.com/rLannes/Rust_get_junctions) was used for recovering junctions from BAM files over a genomic interval. Primer3 and blast v2.16.0 were used to design and validate the primers used for RT-PCR. *Compare_conditions* relies on R for its statistical analyses, including the aod package (version 1.3.3).

### Alignment

RNA-seq reads were trimmed using fastp 0.23.4 (default parameters) and aligned using STAR 2.7.9a. For Fig 4C and Fig 6B, read count was normalized to the total number of uniquely mapped reads scale by a factor of one million. We used FlyBase genome version dmel.r6.48. We identified an issue in the FlyBase annotation for *kl-5* exons 13^th^ and 14^th^. Upon correction, the 13^th^ exon coordinates chrY: 83226-87157 becomes chrY:83228-87157, and the 14^th^ exon coordinates chrY:29254-29392 becomes chrY:29254-29394. This change leads to the change of the codon 3440 from GTC to AGC (Val3440Arg).

## Statistical analysis

Statistical analysis for OmniSplice results was performed using the *Compare_conditions* module of the OmniSplice program. *Compare_conditions* is an add-on module of OmniSplice that provides statistical tests for changes in RNA-seq reads at each exon end, comparing two experimental conditions. For example, a change in the EXON-CLIPPED category relative to the spliced category can be compared between control vs. RNAi conditions. The users define the reference category (e.g. SPLICED category) and the category of interest (e.g. EXON-CLIPPED), then *Compare_conditions* collects RNA-seq reads information from the.junctions files, and applies both a Fisher test and a binomial generalized linear model, defining the reference category as ‘success’, and the category of interest as ‘failure’. Alternatively, *Compare_conditions* can use both a beta-binomial model and a t-test. For each intron, *Compare_conditions* reports (for each exon) the test statistics, the p-values, and the q-value (Benjamini-Hochberg procedure), the delta-psi, the Cohen’s h, the raw read counts per category per condition, and the success/(success + failure) ratio for all conditions. For all analyses including comparison to other software, a gene was marked significant if it has |delta-psi | > 0.1 and adjusted p-value < 0.01 (for MAJIQ we used “ttest-raw_pvalue” < 0.01).

### Back-splicing detection

After our finding that a considerable fraction of the EXON-CLIPPED category likely represents back-splicing events, we generated the back-splicing detection module for OmniSplice. Briefly, the back-splicing module extracts the clipped sequences (from the EXON-CLIPPED category) from OmniSplice output and realigns to the genome using Bowtie2 (2.5.4). For each clipped sequence that aligns to the genome, the best alignment is selected for further analysis. Then, the back-splicing module detects the events where clipped sequences are consistent with back-splicing.

### Fly husbandry and genetics

All fly stocks were raised on standard Bloomington medium at 25°C, and young flies (1- to 3-day-old adults) were used for all experiments.) Control flies were *bam-GAL4* stock. The following fly stocks were used: *U2af38^HMS04505^*(Bloomington *Drosophila* Stock Center [BDSC]:57585), *SRPK^HMS04507^*(BDSC:57587), *bam-GAL4*:*VP16* (BDSC:80579). We used FlyBase (release FB2024_02) to find information on gene sequences/functions, phenotypes, and stocks.

### RT-PCR and RNA isolation

Total RNA was purified from adult testes (100 pairs/sample) by TRIzol (Invitrogen) extraction according to the manufacturer’s instructions. Reverse transcription was performed using the SuperScript IV First-strand synthesis system (Invitrogen) with 3 μg of RNA per genotype and random hexamer primers. PCR was performed using Phusion High Fidelity DNA Polymerase (ThermoScientific). Primer sequence and information are reported in Table S3. PCR products were extracted and sequenced by Plasmidsaurus using their Premium PCR service and sequencing results aligned using ApE and SnapGene.

## Data Availability

The following publicly available RNAseq datasets were used:

Total RNA seq data of the *Drosophila* testis from control, *U2af38RNAi*, and *SRPK^RNAi^* animals (GEO accession: GSE268126) (Fingerhut et al., 2024).

Total RNA seq from the cortex and hippocampus in female wild type and TDP-43 KQ mutant mice (GEO accession: GSE216294) (Necarsulmer et al., 2023).

All other data are provided within the manuscript.

## Acknowledgements

We thank the members of the Yamashita lab and George Bell for discussions and comments on the manuscript. We thank FlyBase, the Bloomington *Drosophila* Stock Center, and the *Drosophila* Genomics Resource Center for reagents and information.

## Author contributions

Conceptualization: RL, JF, YY. Computational analysis: RL. Writing manuscript: RL, YY. Editing Manuscript: RL, YY, JF. Investigation: RL, JF, RLi, RC, AS. Funding acquisition: YY. Supervision: YY

## Supplementary Figures

**Fig S1.**
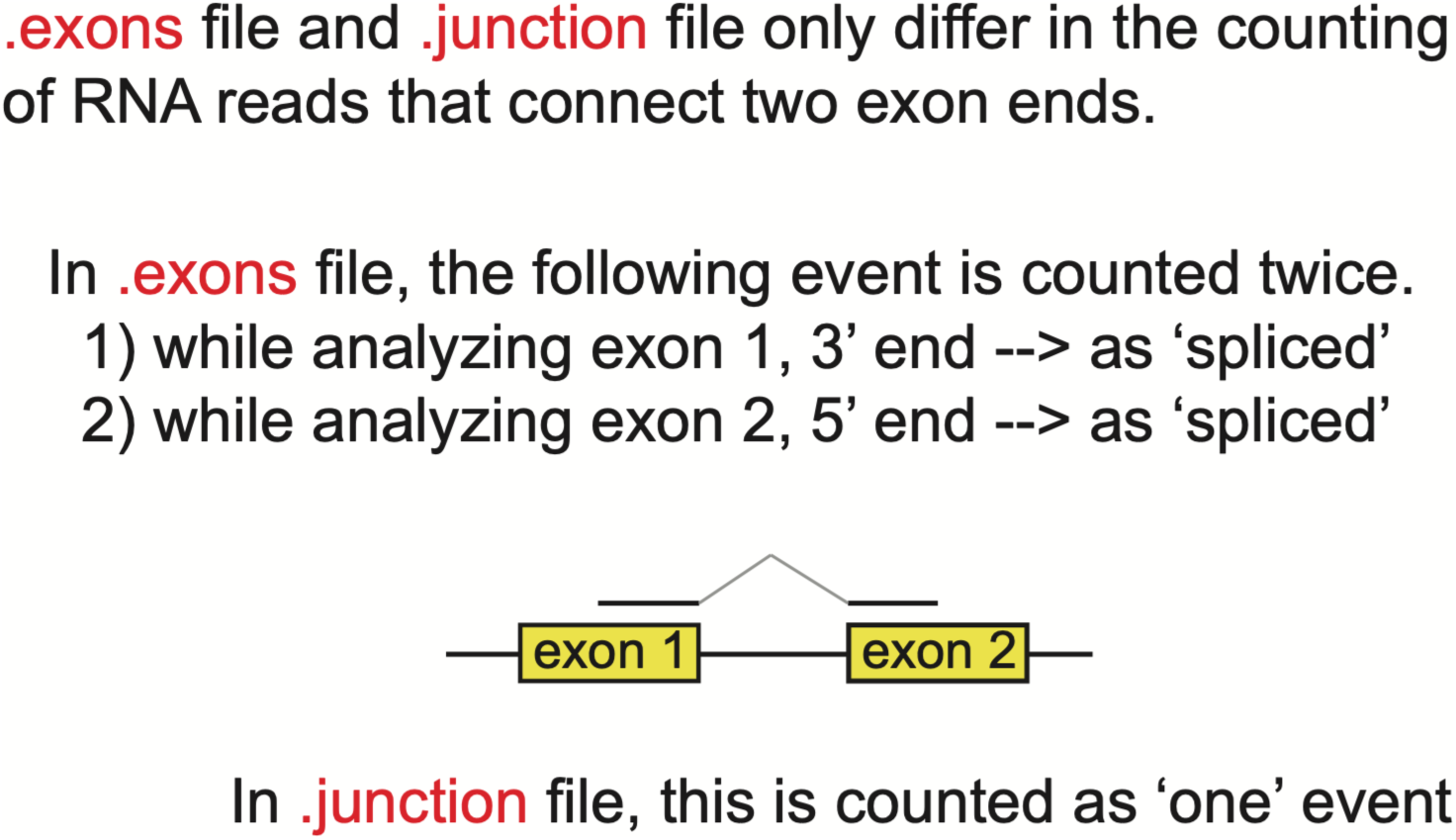
Definition of the.exons and.junctions files.

**Fig S2.**
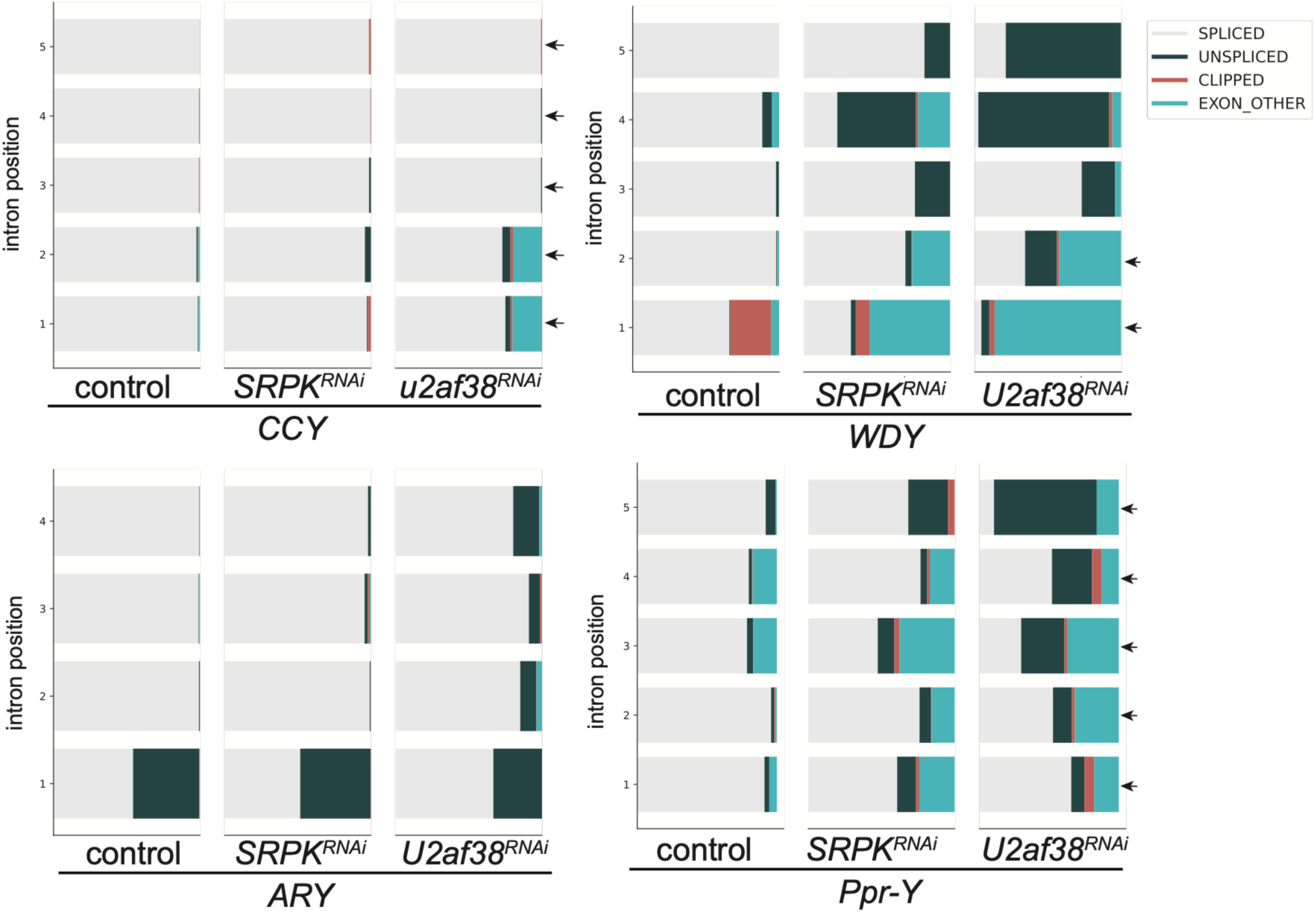
Analysis of splicing status of the Y-linked gigantic genes, *CCY, WDY, ARY* and *Ppr-Y*, in the *Drosophila* splicing mutants. Bar plots showing the splicing status of the Y-linked gigantic genes control, *SRPK^RNAi^* and *U2af38^RNAi^*. Each row represents individual introns, where the proportion of reads from the different categories is shown: SPLICED (light grey), UNSPLICED (dark green), EXON-CLIPPED (orange), EXON-OTHER (blue). Arrows indicate gap-containing introns.

**Fig S3:**
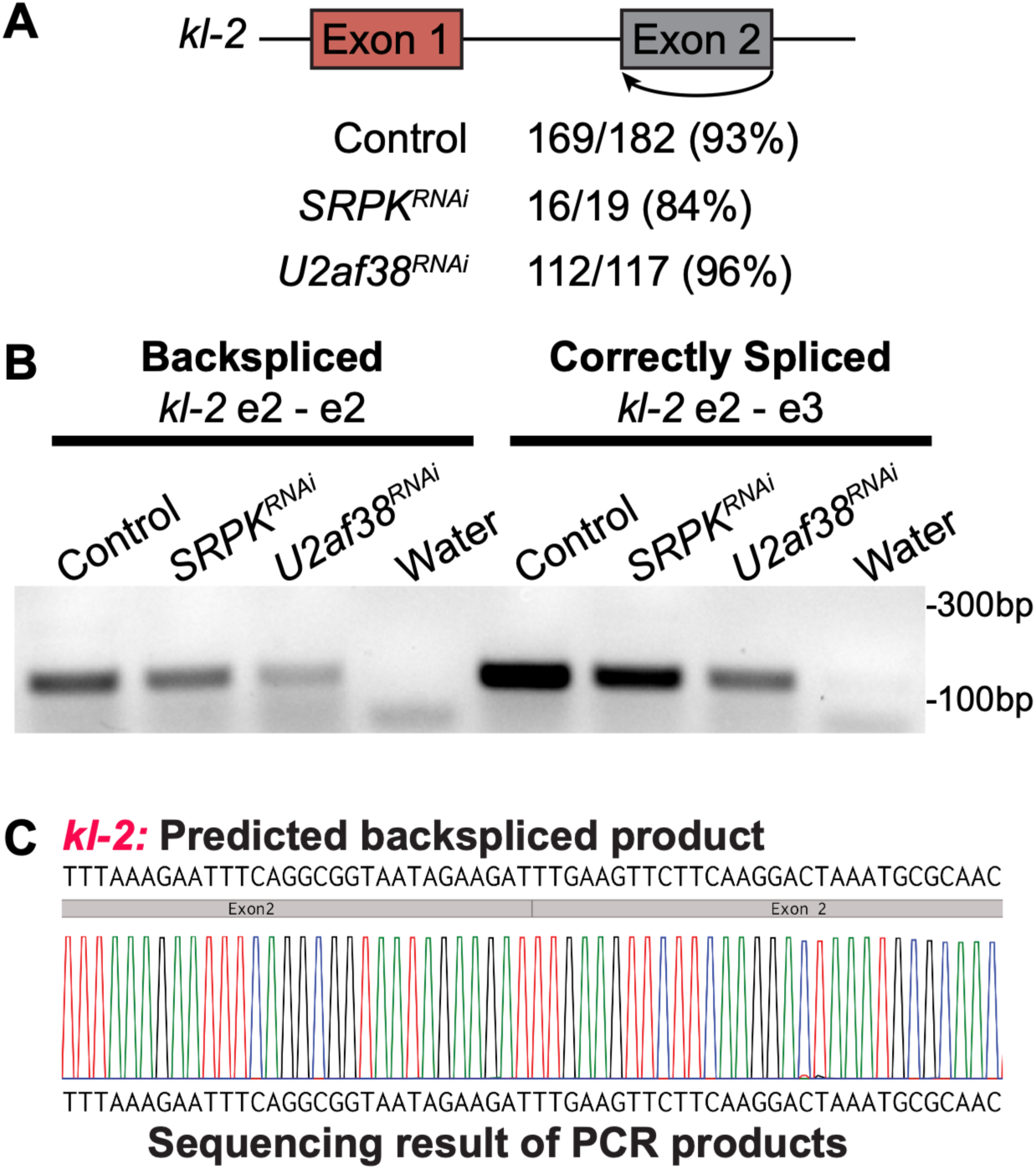
Validation of back-splicing of *kl-2* exon 2. A) Summary of read counts representing back-splicing events/ total ‘EXON-CLIPPED’ reads analyzed by OmniSplice (totaled from all replicates). B) RT-PCR products run on an agarose gel, using primers that are designed to detect back-splicing products (left) and correctly spliced products (right) at the end of *kl-2* exon 2. C) Sequencing results of the PCR product confirming the back-spliced junction.

**Fig S4:**
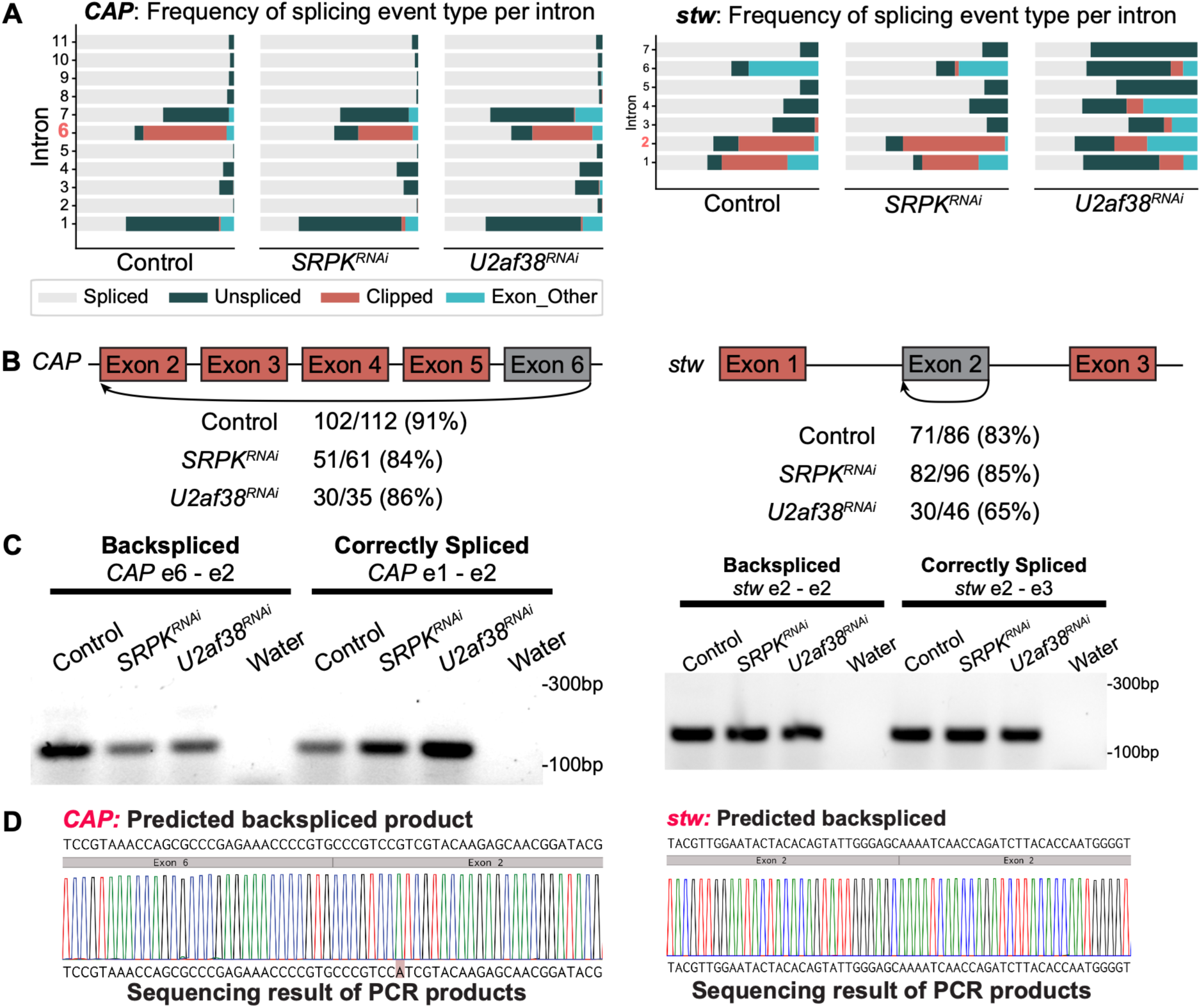
Validation of constitutive back-splicing of *CAP* and *stw* genes. A) Bar plots showing the splicing status detected by OmniSplice in control, *SRPK^RNAi^*and *U2af38^RNAi^* conditions. Each row represents individual introns, where the proportion of reads from the different categories is shown: SPLICED (light grey), UNSPLICED (dark green), EXON-CLIPPED (orange), EXON-OTHER (blue). B) Gene models of *CAP* and *stw* genes, showing detected back-splicing events and raw read counts (totaled from all replicates). Number of back-splicing / total ‘EXON-CLIPPED’ reads. C) RT-PCR products run on an agarose gel, using primers that are designed to detect back-splicing products (left) and correctly spliced products (right) for *CAP* and *stw* genes. D) Sequencing results of the PCR products confirming the back-spliced junction.

**Fig S5:**
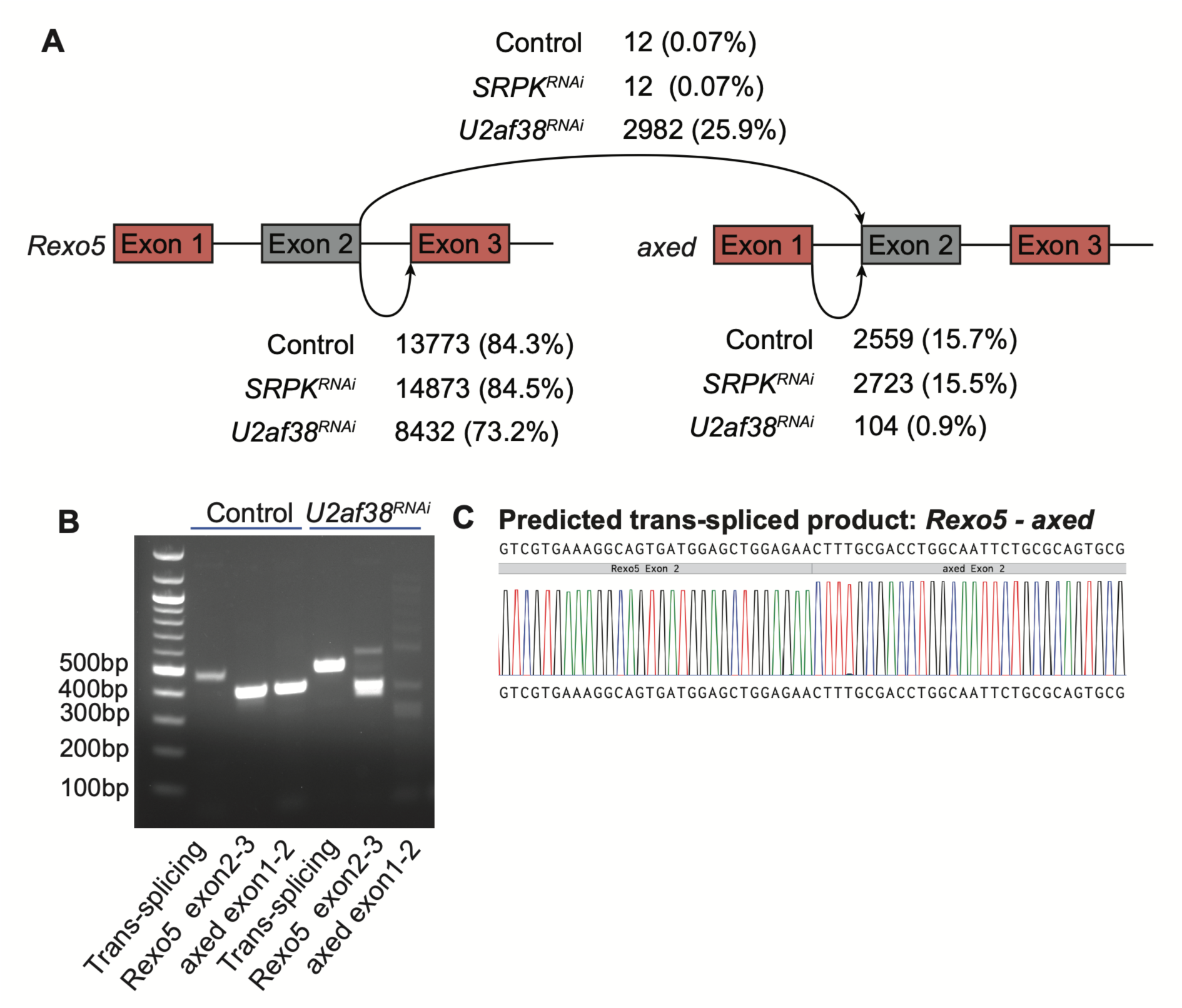
Validation of trans-splicing between *Rexo5* and *axed* in the *U2af38^RNAi^* mutant. A) Gene model of *Rexo5* and *axed*, showing trans-splicing event. B) RT-PCR products run on the agarose gel, using primers that are designed to detect trans-splicing products and correctly spliced products for *Rexo5* and *axed*. C) Sequencing results of the PCR product confirming the trans-splicing junction.

**Fig S6:**
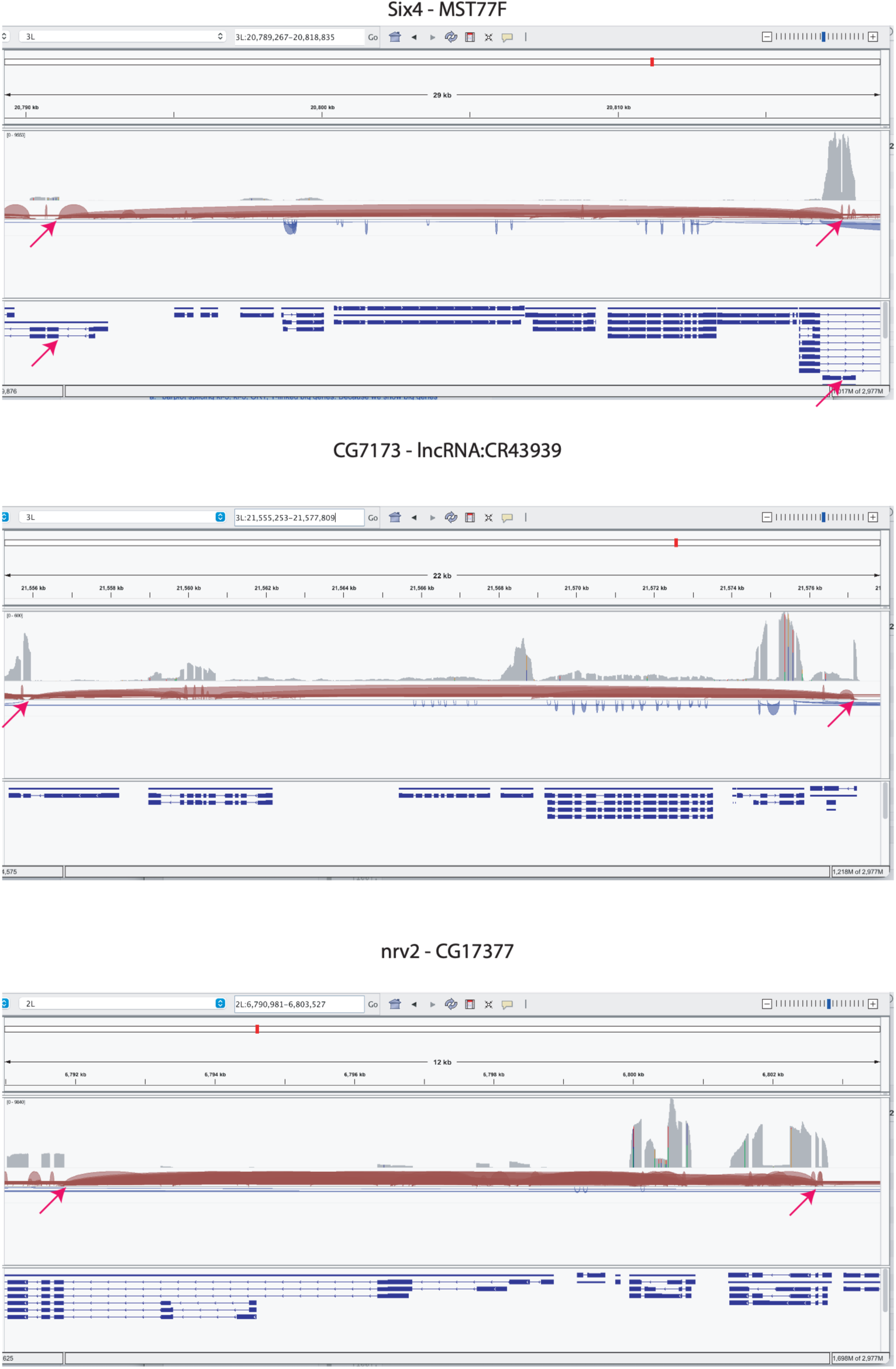
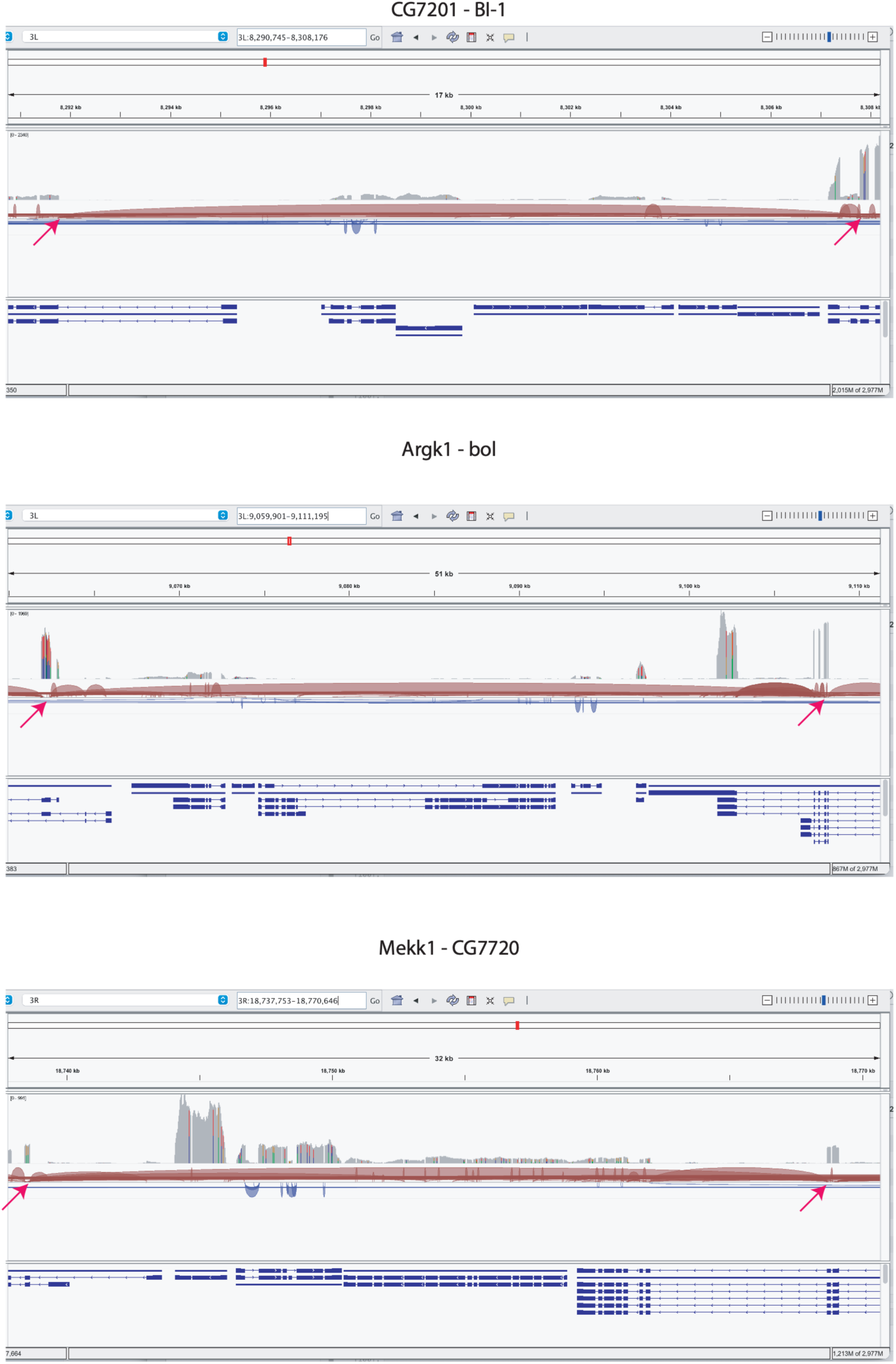
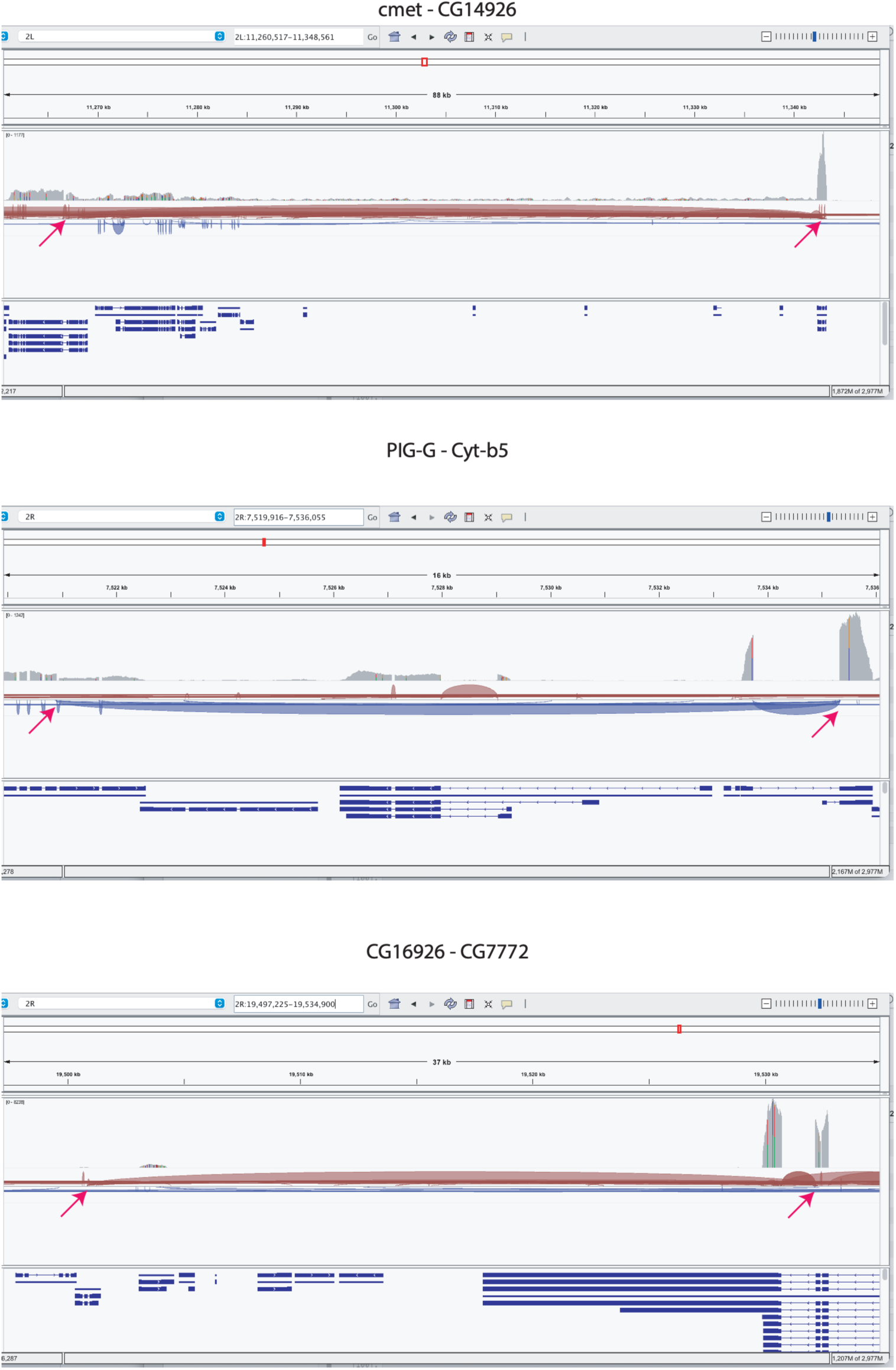
IGV views of representative back-splicing events detected by OmniSplice.

**Fig S7:**
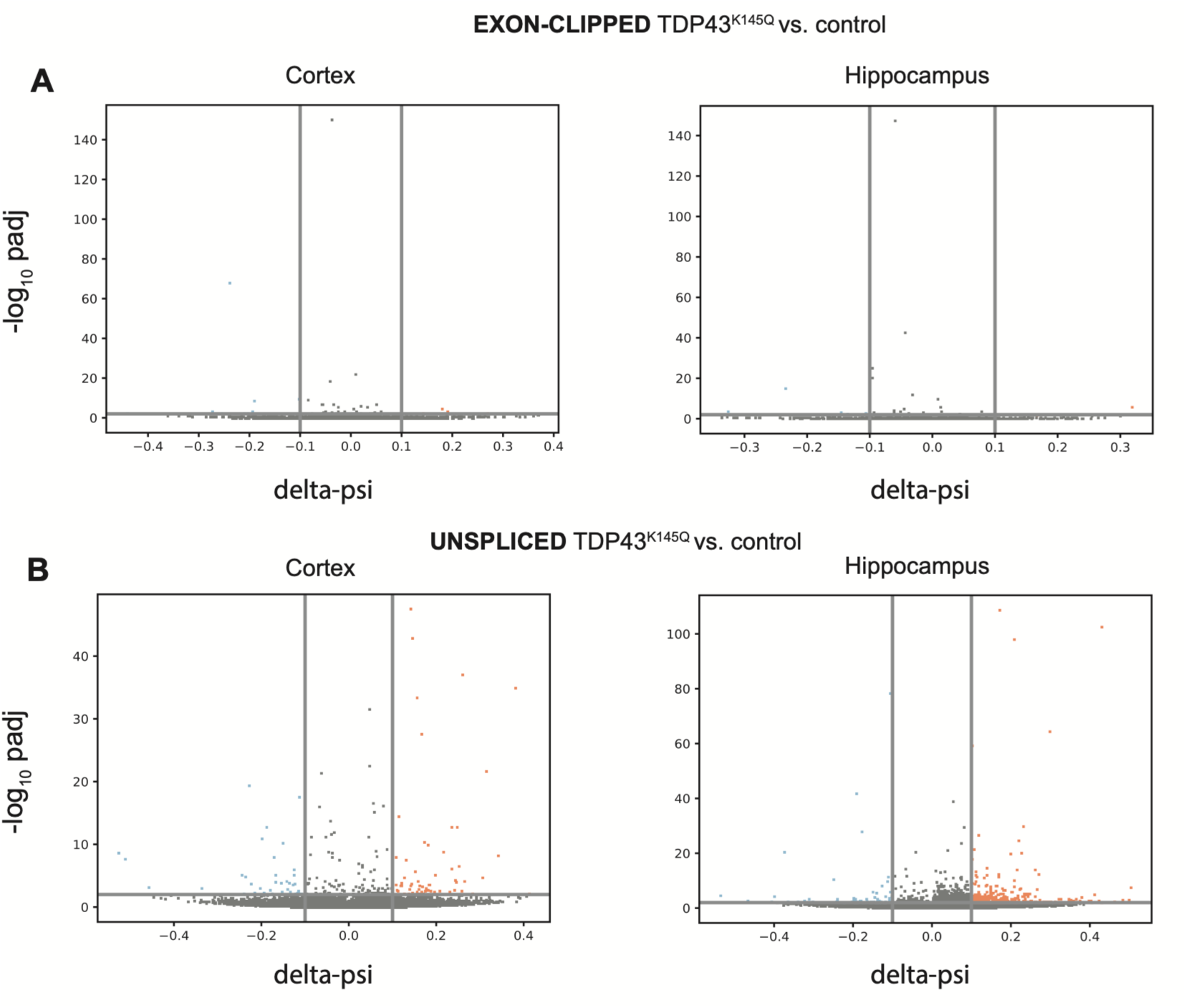
Analysis of EXON-CLIPPED and UNSPLICED categories from TDP43^K145Q^ mutant using OmniSplice. Volcano-style plots for EXON-OTHER category in the cortex (left) and the hippocampus (right) in TDP43^K145Q^ conditions, showing the delta-psi (X-axis) and the adjusted p-value (Benjamini-Hochberg procedure) (Y-axis). The p-value and ratio values were computed using OmniSplice “*compare_conditions*” module, The delta-psi provides the effect size for ratio changes (see materials and methods. A) EXON-CLIPPED, B) UNSPLICED.

## Supplementary Tables (provided as separate excel files)

**Table S1: List of back-splicing events detected by OmniSplice**

**Table S2: List of trans-splicing events detected by OmniSplice**

**Table S3: List of primers used for PCR**

